# Very high extinction risk for *Welwitschia mirabilis* in the northern Namib Desert

**DOI:** 10.1101/2020.05.05.078253

**Authors:** Pierluigi Bombi, Daniele Salvi, Titus Shuuya, Leonardo Vignoli, Theo Wassenaar

## Abstract

One of the most recognisable icon of the Namib Desert is the endemic gymnosperm *Welwitschia mirabilis*. Recent studies indicated that climate change may seriously affect populations in the northern Namibia subrange (Kunene region) but their extinction risk has not yet been assessed. In this study, we apply IUCN criteria to define the extinction risk of welwitschia populations in northern Namibia and assign them to a red list category. We collected field data in the field to estimate relevant parameters for this assessment. We observed 1330 plants clustered in 12 small and isolated stands. The extent of occurrence has a surface of 214.2 km^2^ (i.e. < 5000 km^2^) and the area of occupancy a surface of 56.0 km^2^ (i.e. < 500 km^2^). The quality of habitat is expected to face a reduction of 69.47 % (i.e. > 50 %) as a consequence of climate change predicted in the area. These data indicate a very high extinction risk for welwitschia in northern Kunene and classify these populations as endangered (EN) according to IUCN criteria. Similar assessments for other subranges are prevented by the lack of relevant data, an issue that deserves further research attention. Our results advocate the necessity of a management plan for the species, including measures for mitigating the impact of climate change on isolated populations across its fragmented range.

---

*Welwitschia mirabilis* Hook 1863 is the only member of the family Welwitschiaceae, a relict lineage within gymnosperms endemic to the central and northern Namib Desert. Welwitschia plants are recognized as important components of Namib Desert ecosystems, being exploited by many species of vertebrates and arthropods (Marsh, 1987; Wetsching, 1997; Prendini & Esposito, 2010; Pitzalis et al., 2017). It is one of Namibia’s most iconic species and appears on the National Coat of Arms as a symbol of fortitude and tenacity. In addition, the peculiar morphology and ecology of this plant attracts tourists in the area, contributing to the wellbeing of local human populations.

The distribution of welwitschia is fragmented into four well-separated subranges from the Kuiseb River in Namibia to the Nicolau River north of Namibe in Angola (Kers, 1967; Giess, 1969). The southernmost subrange lays in the central Namib Desert (southern Erongo region). A second subrange is a large area in northern Erongo and southern Kunene regions. A further subrange is represented by a small area in northern Kunene at the border between Namib Desert, Nama Karoo, and savanna biomes. The northernmost subrange occurs in the Namibe province, southwestern Angola. These four areas are separated by about 100 km of unoccupied zones.

Despite the great socio-economic and biological interest on this species, many important aspects of its biology still need to be clarified (Henschel & Seely, 2000). Recent studies suggested that climate change may severely impact *Welwitschia mirabilis* (Bombi, 2018; Bombi et al., 2020), particularly in the northern Namibian sub-range. However, no information is available about the species conservation status, especially for the isolated populations in the remote and wild subrange in northern Kunene. The aim of this study is to assess the extinction risk category (IUCN Standards and Petitions Committee, 2019) of the northern Kunene population based on newly collected field data.

During May 2019, we carried out a field expedition in northern Kunene in order to collect distribution, demographic, and ecological data for *W. mirabilis* in the area. We spent 10 days searching for welwitschia plants by (1) driving at low speed along the available tracks (more than 330 km) and (2) conducting walking transects (more than 65 km in total) across potentially suitable habitats. Our local team members, who have an intimate knowledge of the area from years of herding goats and cattle, targeted our search areas. We inspected about 50 km^2^ in the most suitable area of the known subrange and, based on our field effort and on the local knowledge, we are confident that we have observed in the order of 70-80 % of the actually existing plants.

Overall, we recorded 1330 plants in the area, at elevations between 806 and 991 m above sea level. These plants were clustered in 12 distinct stands, with an inter-stand distance between 1.8 and 30 km. The surface area of each stand varied from 35 to 825,000 m^2^ and, overall, they covered a total surface of about 1.5 km^2^. The number of plants per stand varied between three and ∼400, and the plant density in the stands ranged between 0.1 and 88 plants per 1000 m^2^. All the individuals were thus found in small and relatively isolated subpopulations, resulting in a fragmented pattern.

The extent of occurrence (EOO) is defined as “the area contained within the shortest continuous imaginary boundary which can be drawn to encompass all the known, inferred or projected sites of present occurrence of a taxon, excluding cases of vagrancy” (IUCN, 2001). The aim of this parameter is to quantify the degree to which risks from threatening factors are spread spatially across the geographical distribution of the taxon and it can defined by the minimum convex polygon which contains all the sites of occurrence (IUCN Standards and Petitions Committee, 2019). Following this criterion, the EOO of *W. mirabilis* in northern Kunene is a quadrilateral polygon with a surface of 214.2 km^2^ (Fig. 1).

**FIG. 1.**
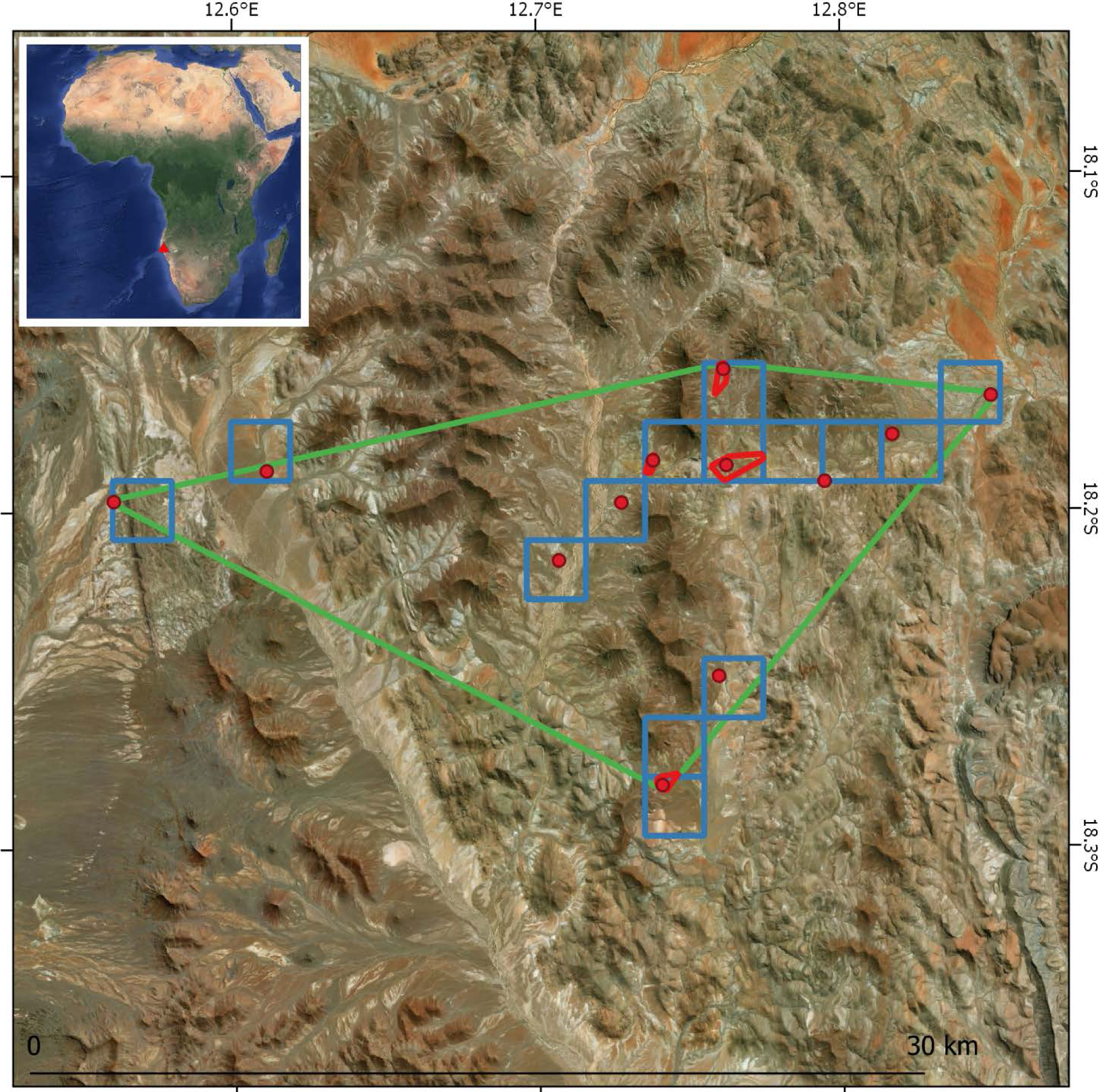
Distribution of welwitschia stands in northern Kunene. Red dots are the centroids of stands and red polygons their boundaries. Blue squares indicate the area of occupancy and the green polygon the extent of occurrence for welwitschia plants. Red triangle in the overview map represents the location of the study area.

The Area of occupancy (AOO) is defined as “the area within its extent of occurrence which is occupied by a taxon, excluding cases of vagrancy” (IUCN, 2012). The twofold role of this metric is to measure the ‘insurance effect’ (i.e. if one taxon occurs in many units of a landscape the probability that it is affected by a threat in all the occupied units is low) and to provide a proxy of the population size, which is generally positively correlated with the number of units occupied. In order to maintain consistency among Red List assessments across taxa, IUCN recommends to estimate AOO using 2 × 2 km grid cells. At this spatial scale, welwitschia plants occupied 14 cells in northern Kunene for a total surface of 56.0 km^2^ (Fig. 1).

To estimate whether the quality of available habitat is likely to change over the time, we used a spatially explicit approach based on ecological niche modeling to predict the climate suitability for the species under recent past and future conditions (see Bombi et al. 2020 for details). To do this, we fitted models on 1000 pseudo-presence/absence points (using EOO as reference) at the spatial resolution of 30 arcsec (about 1 km) in the *R*-based (R Core Team, 2018) *biomod2* Package (Thuiller et al., 2016). Models were fitted using climate data from the WorldClim databank (Hijmans et al., 2005) and then projected under the same climate surfaces as well as under all the available scenarios of future (2050) climate from CMIP5.

Using this approach, we estimated that 246 pixels of the model outputs (93.89 % of the 262 pixels included in the EOO) are facing a negative variation in the quality of habitat, with a mean percentage of suitability reduction of 56.64 % (± 59.73 SD). The reduction of habitat quality is even stronger for the AOO, with 68 pixels (98.55 % of the 69 AOO pixels) facing a suitability reduction. The mean percentage reduction of suitability for pixels in AOO is 72.58 % (± 18.62 SD). Finally, all the 12 stands of welwitschia are experiencing a reduction of climate suitability, with a mean percentage reduction of habitat quality of 69.47 % (± 20.76 SD).

Recent studies suggest that climate change represents the main threat for *W. mirabilis* in northern Kunene (Bombi, 2018; Bombi et al., 2020). Climate change is a global phenomenon, which act at a very large spatial scale, simultaneously influencing populations across the entire distribution of the species but in different ways. Populations can experience negative effects from climate change in some areas of the species range, but positive or no effects in others (Bombi, 2018). Thus, climate change can be considered a threatening factor acting with three different effects in three different locations (sensu IUCN Standards and Petitions Committee, 2019) across the species distribution.

Based on the data we collected and on the results of the modelling analyses, we can apply the IUCN criteria to determine the red list category of welwitschia in northern Kunene. The EOO of the species covers a surface of 214.2 km^2^ (i.e. < 5000 km^2^) and the AOO a surface of 56.0 km^2^ (i.e. < 500 km^2^). Our predictive modeling approach indicates a continuing population reduction based on a decline of habitat quality, both in the past and in the future and where the cause of reduction has not ceased, of 69.47 % (i.e. > 50 %). Climate change is the main threat to the species (i.e. climate change) and can have different effects in terms of magnitude and direction on three locations (i.e. < 5) in the species distribution. These features allow to classify the species as endangered (EN) A4c; B1ab(iii)+2ab(iii) and imply a very high extinction risk for *W. mirabilis* in the northern Kunene.

Unfortunately, no data are available to make comparable assessments for the remaining subranges. The other two Namibian subranges are relatively well known in terms of plant distribution and status, being more easily accessible and closer to main towns than northern Kunene, with some populations being monitored since several years by the Gobabeb Namib Research Institute. Nevertheless, to the best of our knowledge, a complete and accurate dataset on the distribution as well as other relevant information for these populations is not available. In addition, only very general information exists for the Angolan subrange, preventing a species-level assessment of the conservation status. Further research is needed to fill this knowledge gap, especially regarding the virtually unknown Angolan subrange and the real level of isolation between subranges.

In conclusion, our results suggest that this iconic species is particularly sensitive to climate change and there is a real risk of local extinctions in northern Kunene in the next decades. This scenario would be particularly worrisome for the Namib Desert ecosystems as well as for the local human populations. We hope our paper will stimulate the relevant parties at local and international level to set up a management plan for this species, which should take into account measures for mitigating the impact of climate change and other threats in order to guarantee the long-term conservation of *Welwitschia mirabilis*.

## Author contributions

All the authors participated in designing this study, collecting and analysing the data, and writing the paper.

## Acknowledgements

We wish to thank our Himba guides and friends Riatunga Koruhama and Mavekaumba Tjiposa as well as the Okondjombo Community. We are also grateful to the Okondjombo Communal Conservancy; Karen Nott of IRDNC, who facilitated access to the area; Vera De Cauwer of the SCIONA project (EuropeAid/156423/DD/ACT/Multi), for the constructive collaboration; Norbert Juergens of the University of Hamburg, for the helpful information. This study was authorized by the National Commission on Research, Science and Technology (Research Permit RCIV00032018) and financially supported by Mohamed bin Zayed Species Conservation Fund (Project N 182519816) to P. Bombi.

